# Synaptic GABA dysfunction of thalamocortical neurons impairs sleep spindle morphology and fear extinction

**DOI:** 10.64898/2026.05.28.728431

**Authors:** Fumi Katsuki, Meghan C. Bauer, Mitchell J. Vaughn, Victoria A. Lombardi, Ritchie E. Brown, Julie S. Haas, Radhika Basheer, David S. Uygun

## Abstract

The neural mechanisms which control the characteristic waxing and waning shape of sleep spindles and their dysregulation in neuropsychiatric illnesses are unresolved. Recent, sparse research shows post-traumatic stress disorder (PTSD) presents with abnormal spindle morphology and dysfunctional synaptic GABA_A_ receptors in midline thalamic regions. We modeled this GABA_A_ receptor dysfunction by localized CRISPR-Cas9-knockdown in mice. In contrast to a control group with intact GABA_A_ receptors, mice with synaptic GABA_A_ receptor knock-down in thalamocortical (TC) neurons lost the characteristic waxing and waning shape of sleep spindles and exhibited a failure to diminish conditioned contextual fear responses. Our results suggest that abnormally shaped sleep spindles may be a marker of synaptic GABA dysfunction in TC neurons and an indicator of disrupted fear extinction.

**One Sentence Summary:** Abnormal sleep spindle shape reveals underlying inhibitory thalamic dysfunction leading to impaired fear extinction.

## Main Text

Sleep spindles are transient electroencephalographic (EEG) brain waves of light non-rapid-eye-movement (NREM) sleep,^1,2^ named for their characteristic waxing and waning shape.^3^ Typical spindle evaluations quantify their density, measured as the number of spindles per unit of time during NREM sleep, or their amplitude and/or duration, which overlook a key defining feature – their waxing and waning shape. Here we identify a neurobiological substrate for this key feature and show that disruptions of this substrate also leads to alterations in fear processing, a process associated with the pathophysiology of post-traumatic stress disorder (PTSD).

Spindles arise from rhythmic reverberations between the two distinct major neuronal populations of the thalamus^4^, glutamatergic thalamocortical (TC) neurons, and GABAergic thalamic reticular nucleus (TRN) neurons. Spindles are initiated by burst discharges of TRN neurons^5^. However, they are formed by action potential volleys between TRN↔TC neurons.^6^ Burst discharges in TRN neurons produce compound GABA-mediated inhibitory synaptic potentials (IPSPs) in TC neurons that de-inactivate their low-threshold calcium channels, thereby allowing burst discharges in TC neurons following the offset of the IPSP. Burst discharges of TC neurons then excite TRN neurons, restarting the cycle. This reverberation happens at the rhythm of spindles, 10–15 Hz, termed the ‘sigma (σ) band’ in NREM sleep.^7^ Conferred by TC cells to the cortex where spindles are recorded, it defines light NREM sleep. The physiologic mechanisms governing the morphology of canonically defined spindles, i.e. their waxing and waning shape, remain undetermined by biological data. An *in silico* model^8^ suggests that recruitment of additional TRN↔TC neurons and synaptic depression could contribute to waxing and waning,^8^ but these predictions have not been tested *in vivo*.

Although spindle dysregulation occurs in several neuropsychiatric disorders^7,9^, morphology-related parameters may be relevant to PTSD more specifically.^10,11^ Fine grain analysis of spindle abnormalities may shed light on the underlying pathophysiology of PTSD, and lead to better diagnostics or novel therapeutics. Currently, only two studies have evaluated spindle morphology in PTSD and the results are inconsistent. However, both studies show *changes* in spindle quality without a change in spindle density. One reports larger amplitude, plateau shaped spindles in PTSD.^10^ The other shows a relative increase in fast oscillating spindles, but no amplitude change.^12^ Circuit mechanisms which could account for such morphological changes are unclear, but converging lines of evidence suggest thalamic GABAergic dysfunction underlie PTSD symptoms, and waxing/waning morphology may be a readout of this circuit dysfunction: 1) Synaptic GABA_A_ receptors in midline thalamic nuclei that are involved in stress, anxiety, avoidance behaviors, and sleep-wake regulation^13–16^ are structurally altered or possibly lost in PTSD patients^17^. 2) ‘Over consolidation’ of intrusive, stressful memories in PTSD coincide with spindle shape abnormalities.^10^ 3) *In silico*, TRN↔TC “ping-pong” activity shapes waxing and waning.^8^ 4) *In vivo,* spindles are formed by GABAergic signaling in the thalamus which is conferred by TRN→TC connectivity^6^.

Thus, here we used *in vivo* CRISPR-Cas9 knock-down (KD) localized in TC neurons to test whether their synaptic GABA_A_ receptors are necessary for the canonical waxing and waning shape of spindles, and the effect of the KD on fear extinction.

## Results

### TC-α1KD attenuates NREM sleep sigma band power

In adult mouse brains, synaptic GABA_A_ receptors are comprised primarily of two α, two β, and a γ or δ subunit.^18^ γ containing receptors are synaptic and aggregate with α1–3 or α5. In contrast, δ containing receptors are extrasynaptic and aggregate with α4. In TC nuclei of adult mice, synaptic GABA_A_ receptors are exclusively the α1-containing isoform.^19^ To target synaptic receptors, we bred mice expressing Cas9 in vesicular glutamate transporter 2 positive (vGlut2+) neurons, by crossing Vglut2-Cre mice with Rosa26-Lox-stop-lox-Cas9-GFP mice. We termed these offspring Vglut2-Cas9/GFP. We then constructed adeno-associated viral (AAV) vectors to deliver three separate single-guide RNAs (sgRNA) targeting mouse GABRA1, the GABA_A_ receptor α1-subunit gene. We termed this vector AAV5-α1-sgRNA-mCherry (Figure 1A). Each sgRNA is driven by U6 promoter. Separately, the red fluorescent protein mCherry is driven by the human synapsin promoter.

**Figure 1.**
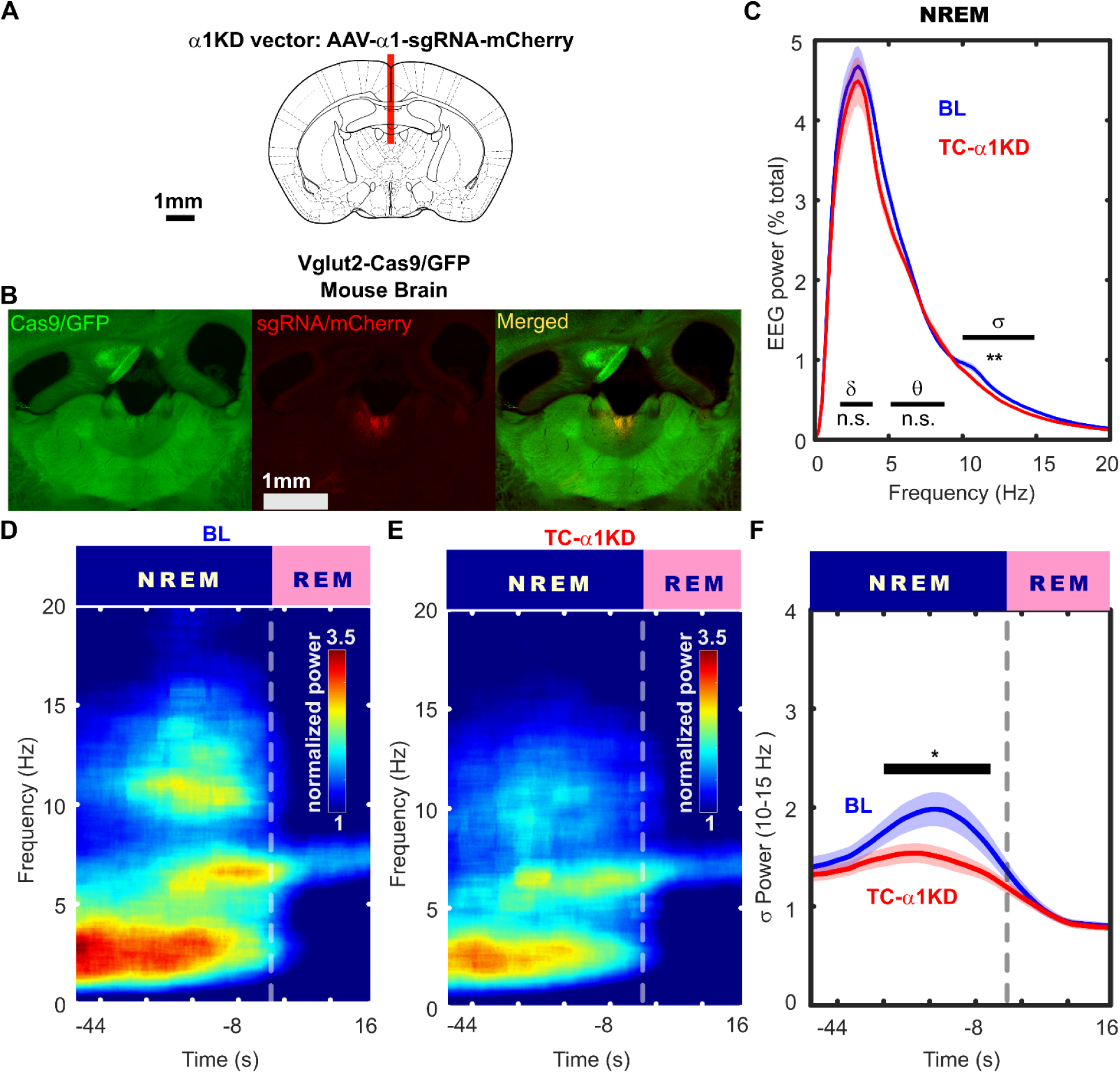
Localized knock-down of synaptic GABA_A_ receptors in thalamocortical (TC) neurons, targeting the paraventricular thalamus (PVT) strongly attenuates NREM sleep sigma (σ) band power (10 – 15Hz) **A.** We designed an AAV vector encoding triple-sgRNA sequences and a fluorescent marker protein, mCherry. To selectively target synaptic GABA_A_ receptors, sgRNAs target the α1 subunit for knock-down (α1KD). α1 subunits are a necessary structure of synaptic GABA_A_ receptors of TC neurons. We injected AAVs in the PVT of Vglut2-Cas9 mice. Together, the sgRNA and Cas9 form a complex that causes α1KD. **B.** Micrographs reveal PVT targeting. GFP, the marker of Cas9 is green. mCherry is red, shown here is our most localized targeting of the PVT, typical expression patterns were more widespread. Yellow shows co-expression, where α1KD occurs. **C.** The power spectrum of NREM sleep EEG in baseline (BL) vs α1KD conditions in the same mice (repeated-measures), revealed a significant reduction in sigma band power (σ; 10 – 15Hz). No change was found in other defined frequency bands, delta (δ) and theta (θ) shown here. **D.** Under baseline (BL) conditions, sigma power surge which is characteristic of wild-type mice is clear in the period of NREM immediately preceding a transition to REM sleep. The sigma power surge is the area of high energy between 10 – 15Hz at approximately-40 and-2 seconds that rapidly disappears in REM. **E.** The sigma power surge is reduced when sleep was recorded after α1KD. **F.** Comparing the sigma power surge in BL vs a1KD revealed a significant reduction following a1KD. n.s. indicates not significant, *p<0.05, **p<0.01. N = 14.

We recorded 24h EEG from frontal cortex, first during baseline (BL) before AAV-α1-sgRNA-mCherry injection, then after injection into the paraventricular thalamus (PVT, Figure 1B), targeting TC nuclei (TC-α1KD). Percentage of time spent in wakefulness, NREM and REM sleep did not change in light or dark periods (Figure S1A). Time-weighted bouts also did not change (Figure S1C-E).

In contrast to BL records, TC-α1KD mice had diminished NREM sigma (σ, 10–15 Hz) power in both light (Figure 1C; p = 0.0033, two-tailed paired t-test;-13.27±3.47% change) and dark periods (p = 0.0003, two-tailed paired t-test;-10.92 ± 2.17% change) and the same frequency band in waking, termed alpha (α, 10–15 Hz) power, during the dark (Figure S2A; p = 0.0002, two-tailed paired t-test;-9.6 ± 1.79% change), but not the light period (Figure S2B). We found no changes to the power spectrum of REM sleep in darkness (Figure S2C), but attenuated REM sleep delta power in the light period (Figure S2D; p = 0.018, two-tailed paired t-test;-14.58 ± 5.1% change). Only NREM sigma power was reduced in both light (Figure 1C) and dark periods (Figure S3A) and was therefore the most robust sleep-wake specific power spectral effect following TC-α1KD. Other bands were unchanged.

In mice, spindles coincide with so-called ‘sigma power surges’, during ∼30s of NREM immediately before REM transitions.^20,21^ In BL, sigma power surges were present (Figure 1D). Following TC-α1KD, sigma power surges were almost abolished (Figure 1E) [σ 10–15 Hz: t (13) =-2.45, p = 0.03; two-tailed paired t-test] (Figure 1F). NREM preceding a transition to wakefulness also produced attenuated sigma power after α1KD (Figure S3B).

### Characterizing sleep spindle morphology in TC-α1KD mice

After finding reduced NREM sigma power, we predicted spindles would be affected.^22^ We were initially surprised to find no reduction in NREM spindle density (N/min). BL was 6.23±0.19 vs α1KD with 6.19±0.29 [t (13) = 0.1, p = 0.92]. Spindle frequency was also unchanged, BL vs TC-α1KD at 12.00±0.03Hz alike. However, spindle duration was markedly reduced following TC-α1KD at 1.82±0.05s vs 1.97±0.03s in BL [*t* (13) = 3.39, *p* = 0.005]. However, in-depth analysis of spindle morphology revealed abnormalities following TC-α1KD. For this, we aligned spindles from each mouse by peaks, and averaged the aligned spindles, canceling asynchronous signals.

This revealed clear waxing/waning profiles in BL spindles (Figure 2A); each peak was larger than the last until the maximum; then each peak was smaller than the last until termination.

**Figure 2.**
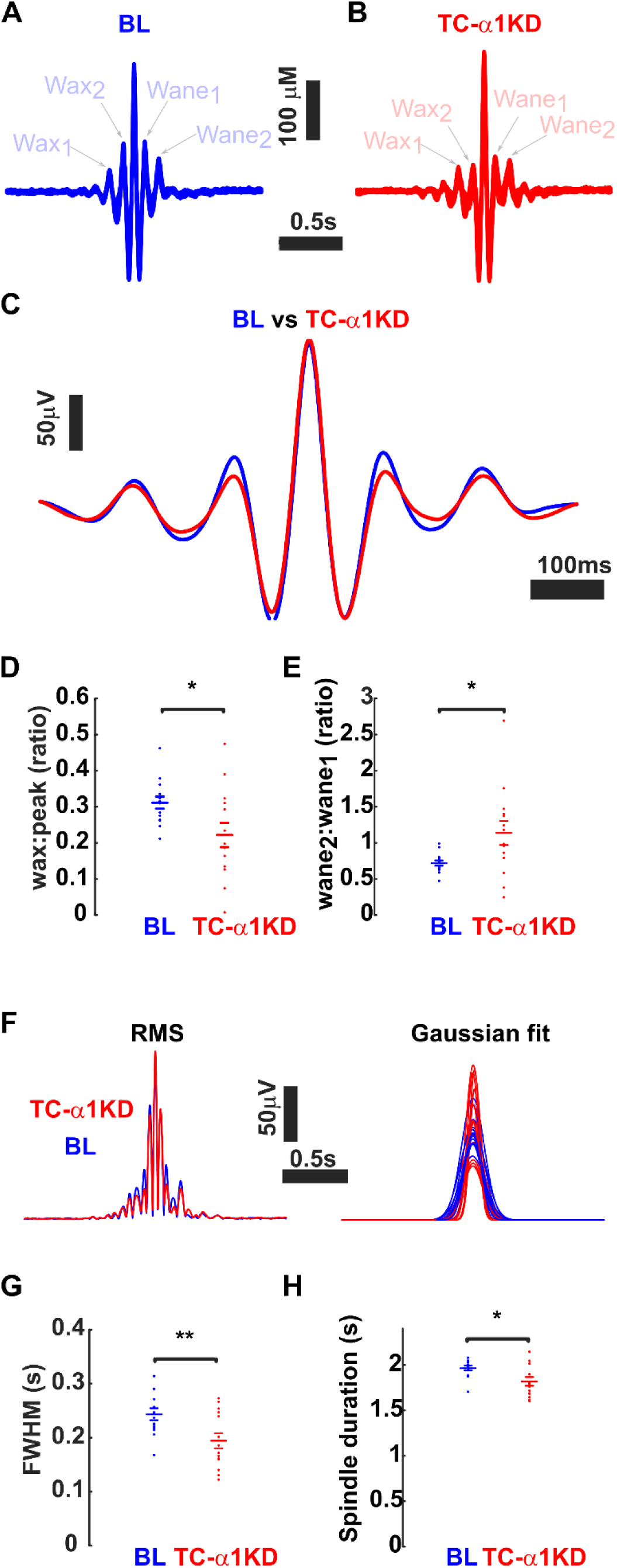
Sleep spindle morphology is abnormal after TC-α1KD. **A.** Averaged sleep spindles from baseline (BL, blue) records have a clear canonical waxing and waning shape. **B.** After CRISPR knock-down of GABA_A_ receptor α1 subunits from thalamocortical neurons (TC-α1KD; red) in the same mice, smaller peaks flanking the maximal central peak have lost the waxing and waning profile. **C.** Compared to BL (blue) sleep spindles, overall power in both the waxing and waning periods is reduced after TC-α1KD (red). **D.** Compared to their BL (blue) records, sleep spindles have a reduced wax to peak ratio after TC-α1KD (red), revealing a larger discrepancy between the waxing part of the spindle and its total amplitude demonstrating a sharper, abrupt, rise than in normal spindles. **E.** Compared to their BL (blue) records, sleep spindles have an elevated wane_2_ to wane_1_ ratio after TC-α1KD (red) revealing a reduced waning trajectory, demonstrating an abnormal termination of sleep spindles. **F.** Root-mean-squared transforms of the spindles were used to rectify the signal to a fit Gaussian function to quantify the shape of the sleep spindles. Visually this reveals a reduction in waxing and waning of the spindles after TC-α1KD. **G.** Full width at half maximum (FWHM) was derived from the Gaussian fit of the sleep spindles and revealed that compared to their BL (blue) records, sleep spindles have reduced FWHM after TC-α1KD (red). **H.** Compared to their BL (blue) records, sleep spindle durations were shorter after TC-α1KD (red). *p<0.05, **p<0.01. N = 14.

Qualitatively, TC-α1KD spindles appeared sharper and the gradual waxing/waning shape (Figure 2B,C) was lost; each peak had a similar amplitude except the central peak.

To quantify spindle shape, we looked at the prominence of waxing and waning by computing the ratio wax_2_:spindle peak, and wane_1_:spindle peak. We found significantly reduced waxing following TC-α1KD (Figure 2D) [t (13) 2.28, p = 0.04], but not waning [t (13) = 1.83, p = 0.09]. We further evaluated the canonical waxing and waning spindle shape, i.e. the crescendo and decrescendo, by analyzing the ratio of the first waxing peak (wax_1_; Figure 2A&B) over the second waxing peak (wax_2;_ Figure 2A&B) and decrescendo as the ratio of wane_2_ over wane_1_. Thus, lower values of either ratio represent the crescendo of the waxing phase, and the decrescendo of the waning phase – canonical in defining spindles. Values near or greater than 1 are divergent from this characteristic. Compared with the BL, wane-ratio (mean ± SEM = 0.54 ± 0.03) was significantly elevated above one [t (12) = 2.52, p = 0.03] following TC-α1KD (mean ± SEM = 1.68 ± 1.01)(Figure 2E). We also found significantly more variance here [F (13) = 0.048, p = 2.9^-6^] following TC-α1KD suggesting spindle tuning disruption.

Interestingly, there was significantly more variance in wax-ratios [F (13) = 0.0010, p = 5.9^-17^] following TC-α1KD, despite a lack of mean change [t (13) = 1.1277, p = 0.28]. Finally, we used RMS-transformed spindles (Figure 2F, left) to fit a Gaussian function (Figure 2F, right), and measured full widths and half maxima (FWHM; Figure 2G). After TC-α1KD, spindles became sharper and more spike-like, as determined by significantly reduced FWHMs (Figure 2H) [t (13) = 3.07, p = 0.0089]. This fitting reveals aberrant waxing and waning morphology of spindles in the KD.

### TC-α1KD impairs fear extinction

PTSD patients have reduced thalamic synaptic GABA_A_ receptor function^17^ and abnormalities in spindles.^10,12^ We hypothesized that reduced thalamic synaptic GABA_A_ receptor function, in mice, simultaneously underlies both altered spindle morphology and disrupted fear extinction, a key symptom of PTSD.^23^ To assess this, we evaluated fear extinction following contextual fear conditioning. We measured time spent moving, interpreting freezing as fearful behavior. To ensure mice were conditioned to aversive training only once as naïve animals, we switched to a between-groups design, using a separate control-AAV injected group. We labelled the control group mice TC-α3KD; they received AAVs delivering sgRNAs that target GABA_A_ receptor α3 subunits, which are not expressed in TC neurons^19^, to control for genomic damage (N = 7). Mice were given two 1.5mA, 5V foot shocks, each two seconds long and separated by 1 minute.

Immediately before the aversive stimuli, both TC-α1KD mice and control mice spent similar amounts of time moving (Figure 3A). On the first visit back to the conditioned context, a day after the aversive stimuli, both groups showed a similar level of fear response and moved for the same cumulative time during the session, indicating that both groups acquired and recalled fear memory in the conditioned context. However, during subsequent days repeatedly visiting the conditioned context without additional foot shocks, control mice clearly returned towards less freezing (Figure 3A, Blue line), indicating fear extinction. In contrast, TC-α1KD mice did not show a return toward baseline levels of movement in the fear-conditioned context. We interpret this as a deficit in contextual fear extinction. ANCOVA revealed a statistically significant difference between the two groups, TC-α1KD mice vs controls, in response to fear extinction trials [F (1, 66) = 7.58, p = 0.0076]. A post-hoc Tukey’s honestly significant difference procedure was used to determine a significant difference between the intercepts of controls vs TC-α1KC mice.

**Figure 3.**
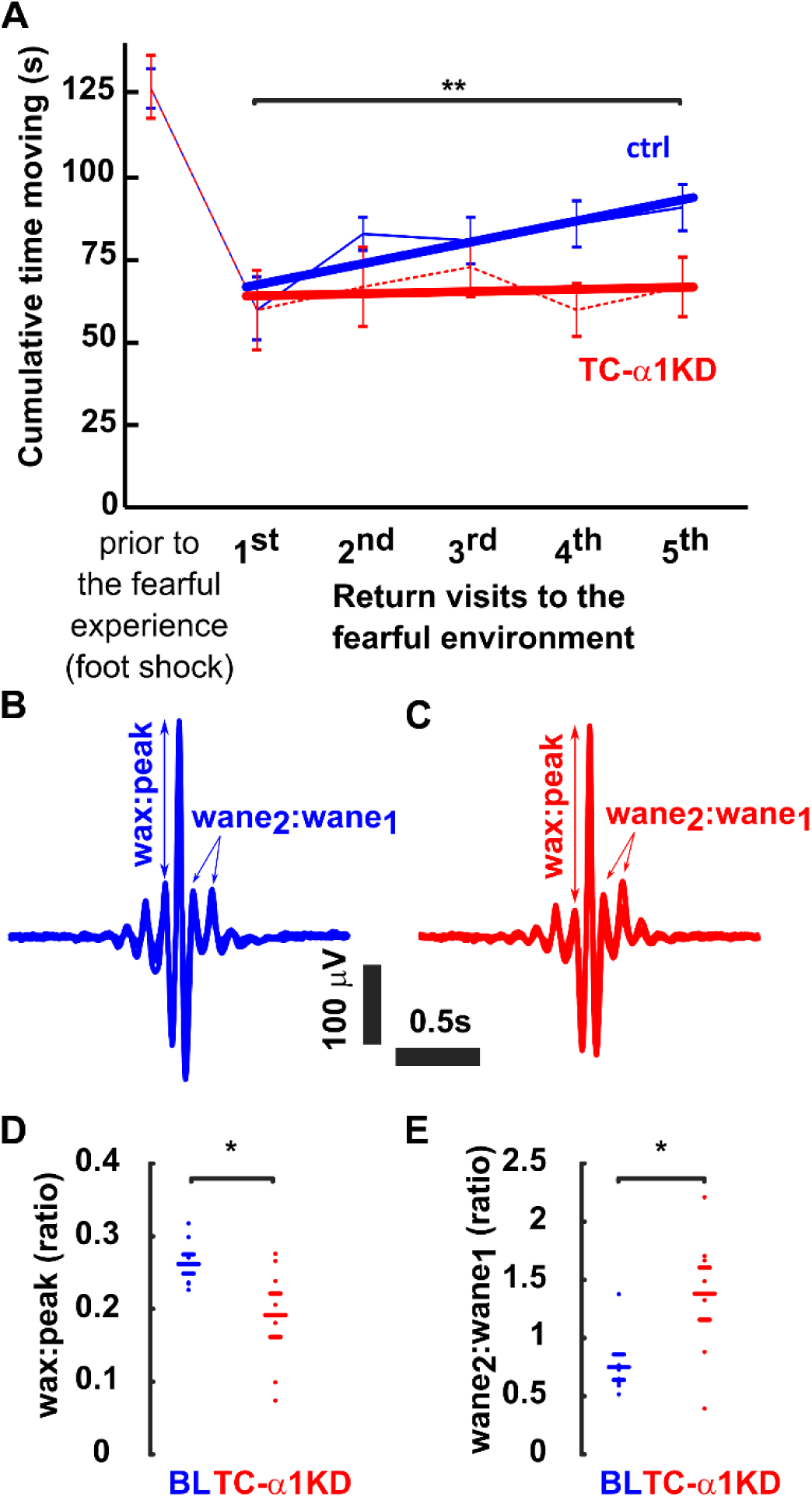
Mice with synaptic GABA_A_ receptors removed from their thalamocortical neurons have diminished recovery from a fearful experience and have abnormal sleep spindle morphology. **A.** Control mice (ctrl; blue) that received a sham KD by AAV bearing sgRNAs against a GABA_A_ receptor subunit not expressed within TC relay neurons of mice (α3) exhibited less time moving in an environment in which they were conditioned to expect a foot shock, on subsequent visits to the fear conditioned environment they became more mobile. By contrast, mice with TC-α1KD (red) reduced their movement to essentially the same amount as control mice. However, TC-α1KD mice continued to exhibit the same low mobility in subsequent visits to the environment, suggesting a failure to recover from the fear of the environment. Thin lines connect mean values; thick lines represent ANCOVA fit lines. **B.** Averaged sleep spindles from the control mice (blue) used in fear conditioning experiments had a clear canonical waxing and waning shape. **C.** Averaged sleep spindles from TC-α1KD mice (red) used in fear conditioning experiments had smaller peaks flanking the maximal central peak have lost the waxing and waning profile. **D.** Compared to the control mice (blue) used in fear conditioning experiments, sleep spindles have a reduced wax to peak ratio in mice with TC-α1KD (red), revealing a larger discrepancy between the waxing part of the spindle and its total amplitude demonstrating a sharper, abrupt, rise than in normal spindles. **E.** Compared to the control mice (blue) used in fear conditioning experiments, sleep spindles have an elevated wax_2_ to wax_1_ ratio after TC-α1KD (red) revealing a reduced waning trajectory, demonstrating an abnormal termination of sleep spindles. \**p*<0.05, \*\**p*<0.001. N = 7/group.

Next, we evaluated sleep spindle morphology (Figure 3B&C) and found the wax-prominence was significantly reduced in the TC-α1KD mice vs controls [Figure 3D; t (12) = 2.1536, p = 0.0262], whereas the wane-prominence was unchanged [t (6) = 0.496, p = 0.32], consistent with our first (within-groups) cohort of mice (Figure 2). The waxing crescendo was reduced to trend level in the TC-α1KD mice vs controls [t (12) = 1.7799, p = 0.0502], but the waning decrescendo was significantly reduced in the TC-α1KD mice vs controls [Figure 3E; t (12) = 2.5342, p = 0.0131], also consistent with our first (within-groups) cohort. The percentage of time spent in wakefulness, NREM sleep, and REM sleep, or bout profiles were not different between the two groups.

### TC-α1KD in vitro verification

To verify TC-α1KD and identify changes to synaptic responses, we performed whole-cell patch-clamp recordings of PVT neurons in acute brain slices prepared from vGluT2-Cas9 mice injected with AAV5-α1-sgRNA-mCherry (Fig. 4 A-C). Control recordings were from uninjected age-matched mice. Recordings were made with CNQX (20 µM), D-APV (50 µM) to block excitatory postsynaptic events, in a high-chloride internal solution internal solution to raise E_Cl_ and amplify inhibitory currents. We recorded 50s of sIPSCs at V_h_ =-70 mV per cell (Fig 4D). We verified that observed sIPSCs were GABAergic by antagonism with bicuculline (Fig 4D, middle traces). IPSCs amplitude from sgRNA-injected animals were not different (Fig. 4E; uninjected: mean 22.4 ± 4.1 pA, n = 11 cells from 2 animals; injected: 19.1 ± 3.4 pA, n = 15 cells from 3 animals, p =0.57, Wilcoxon rank-sum test). However, sIPSC frequency was significantly lower (Fig. 4F; uninjected: mean 5.7 ± 1.0 Hz, n = 11 cells from 2 animals; injected 2.2 ± 0.8 Hz, n = 15 cells from 3 animals; p = 0.005, Wilcoxon rank-sum test). sIPSCs Decay time constants from injected vs uninjected animals were not different (τ_fast_: 9.9 ± 2.2 ms uninjected, 11.8 ± 3.5 ms, p = 0.69; τ_slow_: 39.4 ± 12.1 ms uninjected, 24.7 ± 6.1 ms for injected, p = 0.27, student’s t-test). Intrinsic properties were also not different: input resistance was 470.16 ± 56.8 MΩ for uninjected vs 429.2 ± 34.0 MΩ for injected animals (p = 0.51, student’s t-test); holding currents were-200.9 ± 61.5 pA for uninjected animals and-202.3 ± 43.0 MΩ for injected animals (p = 0.98, student’s t-test) and probability of rebound bursts upon release from 30 mV hyperpolarization was also unchanged (1 for both).

**Figure 4.**
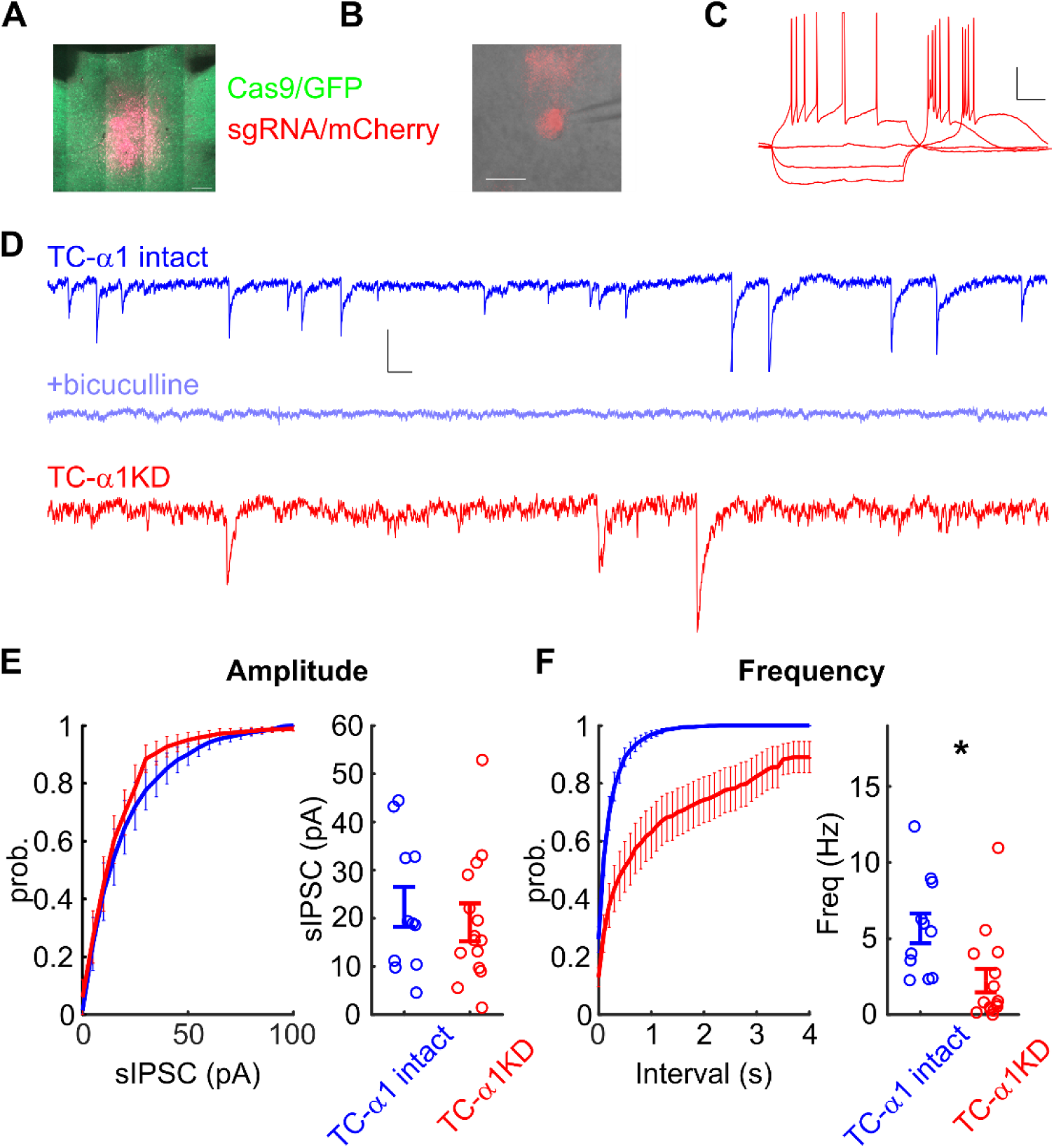
Spontaneous inhibitory postsynaptic currents (sIPSCs) are reduced in frequency by sgRNA knockdown. **A.** Confocal image of injection site. Scale bar 200 µm. **B.** mCherry+ neuron in A. Scale bar 25 µm **C)** Response of a TC-α1KD neuron (sgRNA-mCherry+Cas9-GFP) within the PVT in current-clamp mode to 500-ms pulses of positive and negative currents. V_rest_ =-70.3 mV. Scale bar 20 mV, 100 ms. PVT neurons exhibit rebound bursts upon release from hyperpolarization. **D.** Example traces of whole-cell currents in voltage clamp (V_h_ =-70 mV) with inward sIPSCs from uninjected animals (top), in bicuculline (middle), and from an mCherry+ neuron from sgRNA-injected animals (bottom). Scale bar 10 pA, 100 ms. **E.** Amplitudes of sIPSCs from uninjected (black) and sgRNA-injected animals (red). Mean amplitudes were not different (uninjected: 22.4 ± 4.1 pA, n = 11 cells from 2 animals; injected: 18.0 ± 3.4 pA, n = 15 cells from 3 animals, p = 0.57 (Wilcoxon rank-sum test). **F)** Frequency of sIPSCs in injected animals was significantly lower (uninjected: 5.7 ± 1.0 Hz, n = 11 cells from 2 animals; injected: 2.2 ± 0.8 Hz, n = 15 cells from 3 animals; p = 0.005 (Wilcoxon rank-sum test).

### Histology

We mapped sgRNA-mCherry/Gas9-GFP across multiple nominally distinct thalamic nuclei, by visual comparison with an atlas (Supp Table 1) ^24^ to evaluate the distribution of TC-αKD (Figure 5). One brain from the repeated-measures cohort and one from the TC-α1KD fear conditioning cohort died before we could perfuse them and image their brains. We did not exclude their EEG data, since mCherry was in all the TC-α1KD brains we did image. A single TC-α3KD brain was mCherry–, but we included the *in vivo* data from that mouse because, as a control, it received all procedures.

**Figure 5.**
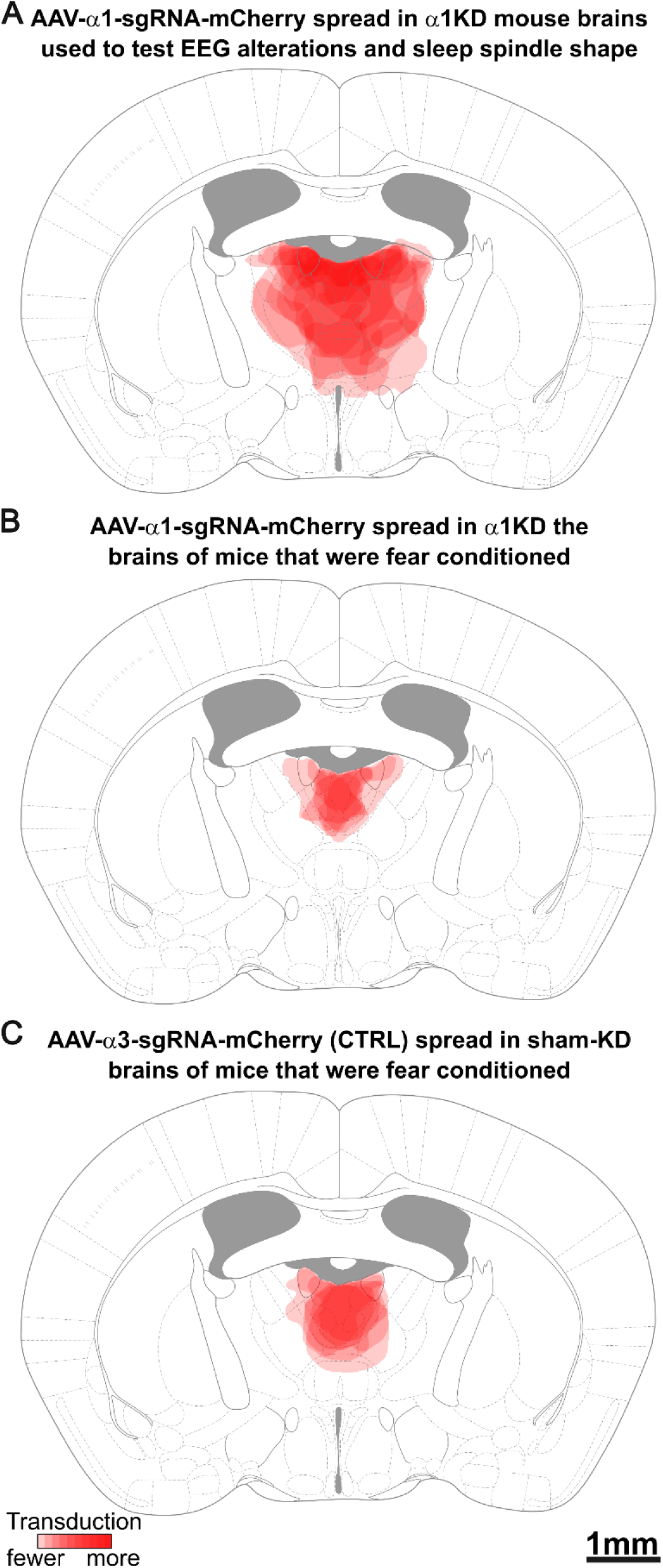
Mapping of AAV distribution patterns in mice after TC-α1KD or TC-α3KD (CTRL). **A.** Composite schematic showing the boundaries of midline injections of AAV-α1-mCherry injections into the midline paraventricular thalamus in 13 of the 14 mice used in the repeated-measures experiments (BL vs KD), with darker red indicating more frequent transduction of the region. Transduced neurons were found more widely beyond the PVT, but were largely restricted to the relay nuclei of the thalamus. **B.** Composite schematic showing the boundaries of midline injections of AAV-α1-mCherry injections into the midline PVT in six of the seven mice used in the between-subjects, fear conditioning experiments, with darker red indicating more frequent transduction of the region. Transduced neurons were found more widely beyond the PVT but were restricted to the relay nuclei of the thalamus. **C.** Composite schematic showing the boundaries of midline injections of AAV-α3-mCherry (CTRL) injections into the midline PVT in six of the seven mice used in the between-subjects, fear conditioning experiments, with darker red indicating more frequent transduction of the region. Transduced neurons were found more widely beyond the PVT but were restricted to the relay nuclei of the thalamus.

## Discussion

Here we show that synaptic GABA_A_ receptor loss in TC neurons simultaneously causes spindle morphological abnormalities and fear extinction deficits, in mice. Given previous reports of thalamic GABA_A_ receptor deficits in PTSD^17^, our findings suggest that spindle morphology may represent a non-invasive indicator of PTSD pathophysiology and symptoms.

We comprehensively characterized spindle rhythmogenesis and found TC-α1KD reduced NREM sigma power, a proxy for spindles^21,22,25,26^, and morphological abnormalities in the spindles themselves. Most studies have not quantified fine-grained morphology of mouse spindles^22,27,28^, we are first to study waxing and waning specifically. We argue this is an important measure since spindles are characterized by waxing and waning, and this shape may be involved in regulating synaptic plasticity.^29^

We propose GABA_A_ receptors on TC neurons sculpt spindles by synchronizing groups of TRN-TC loops to discharge at the same time. The rhythm is set by TRN neurons forming a reciprocally connected loop with TC neurons, and volleying barrages of action potentials.^4^ However, mechanisms synchronizing populations of loops is not understood. One *in vitro* study^30^, proposed spindle waxing occurs as additional TC loops are recruited, oscillating synchronously. Waning occurs as loops stop participating in synchronous oscillation.^30^ Substantial evidence shows spindles are associated with memory.^31^ Thus, accurately synchronizing TC loops may be involved in sleep-dependent memory consolidation, possibly by integrating information from different primary and higher-order thalamocortical circuits. We postulate that the synchronization of neighboring TC loops for waxing and waning requires fast inhibition of GABA_A_ receptors, for timing, since temporally precise inhibition synchronizes networks.^32–34^ This may be one reason TC neurons are enriched with GABA_A_ receptors that have the fastest decay times, those bearing α1 subunits.^19,35–37^ TC-α1KD results in essentially a loss of amplitude surrounding the central peak of the spindle, indicating attenuated synchronous activity between TC-loops – thus the loss of a properly shaped spindle. Elegant experiments by Rovo et al.^23^ delivered Cre recombinase locally to the thalamus in mice with a floxed GABA_A_ receptor γ2 subunit gene. Thus, locally knocking out synaptic GABA_A_ receptors in TC neurons of adult mice, conceptually very similar to our experiments. While they report no major spindle effects, sleep spindle *morphology* analysis is novel here. We also targeted different thalamic regions.^38^ Consistent with Rovo et al., spindle density (N/min) or cycle frequency was unchanged. Also rebound burst firing, key to sleep spindle generation, was intact which is expected since voltage gated calcium and potassium channels were unperturbed.^4^

Emerging research suggests spindle morphology is altered in PTSD.^10^ In health, spindles are associated with consolidating memories.^39^ Earlier studies focused on the amount of spindles in neuropsychiatric illness. However, there is a shift to looking at quality and timing of spindles.^40,41^ Spindle amount is not reduced in PTSD^10,12^, but spindle abnormalities are apparent.^23^ More work is needed to characterize PTSD spindle abnormalities, since the few studies that exist analyzed them differently. Synaptic GABA_A_ receptor dysfunction in the thalamus was evident in PTSD patients who received a radioactive form of flumazenil to visualize synaptic GABA_A_ receptor binding via PET.^17,42^ Reduced binding was evident in three brain areas, including midline thalamus.^17^ Our results show that reduced synaptic GABA_A_ receptors in midline thalamus diminish spindle duration and morphology.

We found TC-α1KD mice had fear extinction deficits whereas fear acquisition was intact. Impaired fear extinction and recall have been reported in PTSD patients.^43,44^ Fear conditioning and extinction involve multiple nuclei including hippocampus, amygdala, and prefrontal cortex. The PVT and prefrontal cortex have a robust reciprocal connection forming parts of fear memory circuits^45^, with infralimbic (IL) cortex and PVT thought to be critical for fear extinction.^46^ Although the IL is implicated in multiple aspects of fear extinction^47–52^, these include consolidation and retrieval of extinction memory.^53,54^ A study in mice showed enhanced long-term potentiation of mediodorsal thalamic inputs to IL promoted consolidation of extinction.^55^ Another demonstrated that chemogenetic inhibition of PVT neurons impaired extinction retrieval.^46^ Our study did not identify which aspects of extinction require synaptic GABA_A_ receptors of TC neurons. However, based on our results that GABA_A_ receptor dysfunction and spindle morphology abnormalities coupled together, and an established association between memory consolidation and sleep, we postulate that GABA_A_ receptors of TC neurons might be key for consolidation of fear extinction. This adds to a small but growing body of literature linking spindles to PTSD.^11,12,56^ Ours is the first to investigate a causal relationship between TRN→TC circuit dysfunction and 1) spindle morphology abnormalities and 2) fearful behaviors, at the same time. If true in humans, a simple algorithm could provide an objective detector of circuit dysfunction in PTSD. This could be a diagnostic and/or screen, or assay for treatment progress.

## Materials and Methods

### In vivo experiments

#### Mice

To use Clustered Regularly Interspersed Short Palindromic Repeats (CRISPR) mediated Knock Down (KD) thalamocortical (TC) neurons, we bred mice expressing the endonuclease of the CRISPR system, Cas9, within glutamatergic neurons positive for the Vesicular glutamate transporter 2 (VGlut2), which is expressed throughout TC neurons. We achieved this by crossing male Rosa26-lox-stop-lox-Cas9/GFP mice (Jackson Labs strain # 026175) with female Vglut2-Cre (Jackson Labs strain #016963) mice. Mice aged 4 to 8 months of either sex were included in all experiments. Mice were housed in a constant light:dark cycle of 12h:12h; lights on (zeitgeber time 0) at 07:00 local time. Ambient temperatures were kept constant between 26–34 °C, with humidity kept constant at 30–70%. Standard rodent chow and drinking water were provided to mice ad libitum. All experimental procedures were first approved by our Institutional Animal Care and Use Committee at the VA Boston Healthcare System, conforming to National Institute of Health, Veterans Administration & Harvard Medical School ethical guidelines.

#### Adeno-associated viral (AAV) vectors

To selectively abscise the gene of synaptic GABA_A_ receptors of TC neurons, (*Gabra1;* NCBI Reference Sequence: NM_010250.5) we developed AAV-α1-sgRNA-mCherry. To produce this, we modified our custom construct previously used to target the gene of the GABA_A_ subunit of the thalamic reticular nucleus (Gabra3) by replacing the U6-promoter-single-guide-RNA triplicate cassette with sgRNAs complementary to Gabra1. We used an online CRISPR sgRNA identification tool, http://chopchop.cbu.uib.no/ ^57^ for identification of candidate sgRNAs. The final three sgRNAs (Sequences here) were selected manually based on proximity the 5’ end of the coding region of the gene. None of the 3 sgRNAs were determined to bind off-target mouse genomic DNA in silico. For our control AAV vector, used in the between-subjects experiments, we used AAV-α3-sgRNA-mCherry which was previously described in Uygun et al. 2022.^20^ Since TC neurons are devoid of α3 subunits, AAV-α3-sgRNA-mCherry causes DNA damage to a non-expressed gene, to control for genomic DNA damage without an actual KD of receptors. Snapgene was used in designing our plasmid constructs, but actual DNA manipulation was performed commercially by GenScript (New Jersey). Plasmids were validated at GenScript via sequencing. Plasmids were then sent to the University of North Carolina Vector Core for packaging into AAV serotype 5 particles at titers ∼1–4 × 10^12^ particles/ml, as determined by dot-blot analysis.

#### Thalamic AAV microinjections and electrode implantation

Selective CRISPR mediated α1KD within TC neurons depends upon colocalization of GABRA1 specific sgRNAs and Cas9 in cells. For this we used stereotaxic injection of AAVs to locally deliver the sgRNAs; Cas9 was expressed by the transgenic mouse genomes. Surgeries were performed with a Leica stereotaxic apparatus. Repeated measures experiments: We first implanted cannulas (Plastics One; Connecticut; Part# C315G/SPC, cut to length 12mm) and their retainers (Plastics One, Part # C15I/SPC) positioned with the embedded tip 2mm above the PVT (from bregma: AP −0.06 mm, ML ±0.0, DV −1.1). Frontal cortical EEG electrode screws (Pinnacle Technology Inc.; Kansas, United States; Part # 8403; from bregma: AP +1.9 mm, ML ±1.5), a reference electrode screw (from bregma: AP −5.2 mm, ML +0.0) and a ground electrode screw (bregma AP-3 mm, ML = 2.7) were implanted and their attached leads were soldered to a Pinnacle Technolology Inc.; Part # 8201-SS headmount. The EMG electrodes from the headmount were positioned underneath the neck muscle. Dental cement (Keystone industries, Bosworth Fastray; Part # 0921378) was used to fix all the implanted materials in place. After at least a week of post-surgery recovery baseline records of EEG/EMG were collected. Then, AAVs were microinjected targeting the PVT using a 5 μl Hamilton syringe (Part # 87908, Model 75 SN SYR with 33 g cemented needle) and a KD Scientific Legato 130 micropump (1 μl at 0.05 μl/minute) by lowering the needle tip 2mm beyond the implanted cannula (DV-3.1mm). 5 weeks after AAV injections, EEG and EMG were recorded again for KD the condition, and comparison with BL. Between-subjects experiments: The same materials and coordinates were used, however no cannula was implanted. Instead, AAV-α1-sgRNA-mCherry or AAV-α3-sgRNA-mCherry was injected to the PVT either at the time of electrode implantation, or skin was sutured, and the electrode implantation was performed in a subsequent surgery. Surgeries were performed using isoflurane anesthesia (1.5–5% in O2). The depth of anesthesia was determined by the mouse’s breath rate, pedal withdrawal reflex and tail reflex. Meloxicam (5 mg/kg; intraperitoneal) was administered subcutaneously post-surgery and at 22–24h post-surgery for pain relief.

#### Electroencephalogram (EEG)/Electromyogram (EMG) recordings

To determine sleep wake-states wakefulness, NREM and REM sleep and subsequently perform analysis of the state-specific waveforms, we collected EEG and EMG on Pinnacle Technology Inc. 3 channel (2 EEG/1EMG) systems (Part # 8200-K1-SL), with Sirenia Acquisition software. Mice were habituated for over 24 hours to being tethered by pre-amplifiers (Pinnacle Technology Inc. Part # 8202-SL) and 24 hour EEG/EMG records were collected (zeitgeber time 0–24). EEG and EMG data were sampled at 2kHz and amplified 100x, then low-pass filtered to 600 Hz.

#### Sleep-wake state staging & EEG waveform analysis

We automated sleep-wake scoring using Sleep-Deep-Learner^58^, our validated transfer learning based tool which automated sleep-wake scoring after learning to score from manual scores by the investigator. Manual sleep-wake scoring was performed as we have previously described. Briefly we used 4-second epochs and defined each state as such: Wake was characterized as desynchronized, low amplitude EEG paired with muscle tone in the EMG which did not necessarily need to appear to be phasic. NREM sleep was characterized as slow, synchronized, large amplitude, EEG waves, paired with low amplitude EMG, with brief bursts considered twitching. REM sleep was characterized by stereotyped ‘sawtooth’ EEG, a sharp spike in the theta range (5–9 Hz) of an accompanying power spectral density profile aided identification, paired with an essentially flatline EMG signal, sometimes contaminated by a pulse not indicative of skeletal muscle tone. Artifacts labeled as epochs may be wakefulness, but with crosstalk between EEG and EMG signals. For behavioral evaluations of sleep such as the proportions of states or bouts, all epochs including artifacts were used. However, artifacts were removed for analysis of the actual waveforms including power spectral density, time-frequency analyses, and spindle morphology evaluation. Manual scoring was done in Sirenia Sleep, automated sleep-wake scoring and analyses of sleep-wake scored data were performed in either MATLAB, excel or LibreOffice Calc.

For bout analysis, bout durations were binned into discrete intervals, summed, and presented as a percentage of total time in the state. We used this approach having read methods in previous sleep-wake bout analysis in mice.^59^

#### EEG power analysis

Power spectral density of each sleep-wake state was determined by MATLAB’s pWelch function, 4-second Hanning window, overlap 50%. Normalized power was determined by the percentage of each frequency bin per state of the total summed power of the whole 24h signal, as in prior work.^60^ For time-frequency spectrogram color plots of state-transitions, we down-sampled signals to 40 Hz, screened for outliers by zeroing values over 10 standard deviations from the mean. We used the Chronux toolbox multi-taper^61^ (Chronux.org) function set with 5 tapers and a 10 second, 100 ms step, sliding window. We obtained a spectrum for each state-transition per mouse and averaged them. Next, each average spectrum was normalized such that each frequency bin was divided by the corresponding frequency bin from wakefulness.

Normalized spectra were averaged between the mice in their group providing a grand mean spectrogram for the group. Mean power in the sigma range (10–15 Hz) was smoothed (5 s moving average window) and plotted with envelopes representing standard errors.

#### Sleep spindle analysis

We detected sleep spindles during NREM sleep with our spindle detection algorithm as previously described and validated.^22^ Next, all sleep spindles detected within each mouse were aligned by the positive peak, as determined by the maximum value of the sleep spindle. Visual inspection of the spindle was used to determine the major appreciable peaks. We decided two peaks could be discerned within the waxing part of the sleep spindle (labeled wax1 and wax2), and two peaks could be discerned within the waning part of the sleep spindle (labeled wane1 and wane2). To capture the peaks in waxing, we obtained the maximum value between 0.22 s – 0.18 s prior to the spindle peak for wax1; and between 0.1 s – 0.06 s prior to the spindle peak for wax2. To capture the peaks in waning, we obtained the maximum value between 0.1 s – 0.06 s after the spindle peak for defined wane1; and between 0.22 s – 0.18 s after the spindle peak for wane2. The wax-factor was computed as Wax1 divided by Wax2. The wane-factor computed as Wane2 divided by Wane1. The wax prominence was defined as the peak closest to the central peak (wax2) divided by the central peak. The wane prominence was defined as the peak closest to the central peak (wane1) divided by the central peak. Each of the averaged sleep spindles were root-mean-squared using the MATLAB ‘rms’ function and, subsequently, a gaussian function was fitted using the MATLAB “fit” function with the Gauss1 option. The *c* parameter from the gaussian function was used to calculate full widths at half maximum, as 2√2 *ln* 2 *c* to compare sharpness of the gaussian curves between treatments.

Contextual fear conditioning. Mice were placed in a rectangular chamber (7”X7”X10”, W x L x H) with a surface grid (context A) connected to an electrical shocker (Coulbourn Instruments, USA). Mice were then habituated to the box for 3 min followed by application of two successive foot shocks (1.5mA, 5V) each lasting 2 seconds, separated by an interval of 1 min. Following the shock day, 5-days of extinction sessions were performed. At each extinction session, contextual fear response was assessed by placing the mice in the same context as the shock day for 5 min, but without presenting a foot shock. Contextual fear memory was determined based on the freezing behavior displayed by the mouse, averaged across the first 3 min. Freezing behavior was detected by automated tools within the Noldus EthoVision XT14 video tracking system (Noldus Information Technology Inc.).

### In vitro experiments

#### Whole-cell Patch Clamp Recordings

Vglut2-Cas9 mice were anesthetized under isoflurane and mounted on a Kopf digital stereotax. A small craniotomy was made over PVT under aseptic conditions. 500 nL of sgRNA was injected to PVT (relative to bregma: AP-1.06, ML 0, DV – 3.1) at 100 nL/min using a syringe pump (WPI UMP3). Postoperative analgesia was provided by bupronex (0.1 mg/kg). Lidocaine and antibiotic were topically applied to the wound. Injections were performed on ten-week-old mice, and 4 weeks was allowed for expression prior to recording.

Animals were anesthetized under isoflurane and perfused with ice-cold NMDG solutions (in mM: 92 NMDG, 2.5 KCl, 25 NaHCO_3_, 25 glucose, 1.25 NaH_2_PO_4_, 20 HEPES, 2 thiourea, 5 Na-ascorbate, 3 Na-pyruvate, 0.5 CaCl_2_, 10 MgSO_4_). Horizontal brain slices 300 µm thick were cut with a Leica VT1200S vibratome in NMDG, then incubated in HEPES holding solution (in mM: 92 NaCl, 2.5 KCl, 25 NaHCO_3_, 25 glucose, 1.25 NaH_2_PO_4_, 20 HEPES, 2 thiourea, 5 Na-ascorbate, 3 Na-pyruvate, 2 CaCl_2_, 2 MgSO_4_): at 37°C for 20 min and then in ACSF (in mM: 126 NaCl, 3 KCl, 1.25 NaH_2_PO_4_, 2 MgSO_4_, 26 NaHCO_3_, 10 dextrose and 2 CaCl_2_, 315–320 mOsm, saturated with 95% O_2_/5% CO_2_) at room temperature and through recording. The submersion recording chamber was held at 34°C (TC-324B, Warner Instruments). The approximate bath flowrate was 2 ml/mi and the recording chamber held approximately 5 ml solution. PVT was visualized and fluorescent reporters were verified under 4x magnification. PVT neurons were targeted and patched under 40x IR-DIC optics (SliceScope, Scientifica, Uckfield, UK). Fluorescent light was delivered through the objective (CoolLED pE-300).

Electrodes were filled with (in mM): 130 KCl, 5 NaCl, 2 MgCl_2_, 10 HEPES, 0.1 EGTA, 2 NA_2_ATP, 0.5 GTP-tris, 4 MgATP, pH 7.25 with KOH (290 mOsm). Pipette resistances were 5-9 MΩ before compensation; recordings were discarded if holding currents exceeded 300 pA or access resistance exceeded 25 MΩ. sIPSC recordings were made in voltage clamp with V_h_ =-70 mV. Voltage signals were amplified and low-pass filtered at 8 kHz (MultiClamp, Axon Instruments, Molecular Devices, Sunnyvale, CA, USA), digitized at 20 kHz with custom Matlab routines controlling a National Instruments (Austin, TX, USA, USB6221 DAQ board), and data were stored for offline analysis in Matlab (Mathworks, R2018b, Natick, MA, USA).

Offline analysis was performed in Matlab (R2018b, Mathworks). Postsynaptic currents (PSCs) were detected by convolving a template with the current recording. The template was a product of exponentials: 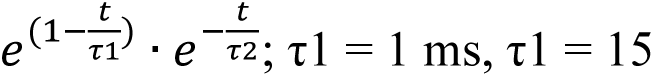 Candidate PSCs were identified by thresholding the derivative of the convolution product, and PSCs over an amplitude threshold were accepted as real PSCs. Both thresholds were manually determined for each cell. Double-exponential fits were made to the decay of the average PSC for each cell.

#### Histology

We performed transcardial perfusion on the mice first using phosphate buffered saline (PBS; 12 ml), then 4% paraformaldehyde (12 ml). We then removed the brains and submerged them in 4% paraformaldehyde for 2 days. We then replaced the paraformaldehyde with 30% sucrose in PBS. Once the brain had sunk in the solution, we sectioned the area of interest to 40 µm using either a microtome or cryostat (Leica Biosystems, Illinois, United States). We mounted the sections slides with coverslips secured by Vectashield Hard Set mounting medium (Part # H-1400, Vector Labor). Micrographs were acquired either by a Zeiss Image2 microscope, using a Hamamatsu Orca R2 camera (C10600) and digitally captured by Stereo Investigator software (MBF Bioscience), or a Vector Polaris SlideScanner with native acquisition software.

To verify successful transduction by AAVs, we visually assessed the presence of mCherry (excitation:emission 590:617) marking sgRNAs, and GFP (excitation:emission 488:509) marking Cas9, throughout the nuclei of the thalamus. We selected sections best corresponding to the brain atlas section at Bregma −1.06 mm, where we had targeted during surgeries and AAV injections. We used a free draw tool in Inkscape to draw around thalamic regions marked by red, within the area marked by green. Note, CRISPR mediated KD is only possible in cells with both red and green, I.E. both sgRNA-mCherry and Cas9-GFP. The transduction area was drawn by hand over micrographs mapped onto an atlas to determine transduced loci. We took a binary approach to labelling transduction whereby a nucleus was said to be transduced if it had mCherry signal in more than half of the region.

## Statistics

Two-tailed paired *t*-tests were used to compare BL vs KD for within subjects experiments. We used two-tailed un-paired *t*-tests in between subjects experiments. ANCOVA was used to compare TC-α1KD vs TC-α3KD (sham) for fear conditioning experiments. Wilcoxon rank-sum test was used in whole-cell patch clamp experiments. All statistics were performed in MATLAB, GraphPad Prism5 or Microsoft Excel.

## List of Supplementary Materials

Materials and Methods Figs. S1 to S3

Table S1 References 57-61

## Funding

VA Biomedical Laboratory and Clinical Science Research and Development Service Career Development Award IK2 BX004905 (DSU)

VA Biomedical Laboratory and Clinical Science Research and Development Service Merit Awards I01 BX001404 (RB)

VA Biomedical Laboratory and Clinical Science Research and Development Service Merit Award I01 BX006105 (RB)

VA Biomedical Laboratory and Clinical Science Research and Development Service Merit Award I01 BX004673 (REB)

VA Biomedical Laboratory and Clinical Science Research and Development Service Merit Award I01 RD001372 (REB).

National Institutes of Health grant T32-HL007901(DSU) National Institutes of Health grant R01-NS119227 (RB) National Institutes of Health grant K01 AG068366 (FK) National Institutes of Health grant NINDS R01 NS128713 (JSH)

## Author contributions

Conceptualization: DSU, REB, RB. Methodology: DSU, FK, VAL, JSH, MCB, MJV.

Investigation: DSU, FK, JSH, MCB, MJV. Visualization: DSU, JSH.

Funding acquisition: DSU. Project administration: DSU. Supervision: DSU, REB, RB. Writing – original draft: DSU.

Writing – review & editing: DSU, FK, JSH, MCB, MJV, REB, RB

## Competing interests

Authors declare that they have no competing interests. DSU is a Research Health Science Specialist, REB, FK, and RB are Research Health Scientists at VA Boston Healthcare System. DSU current address is the University of Warwick, he completed the work on this project while serving as a Research Health Scientist at VA Boston Healthcare System. The contents of this work do not represent the views of the U.S. Department of Veterans Affairs or the United States Government.

## Data, code, and materials availability

All data and code original to this manuscript will be made available by download via the Open Science Framework repository upon formal acceptance to a peer reviewed journal.

---

**Fig S1.**
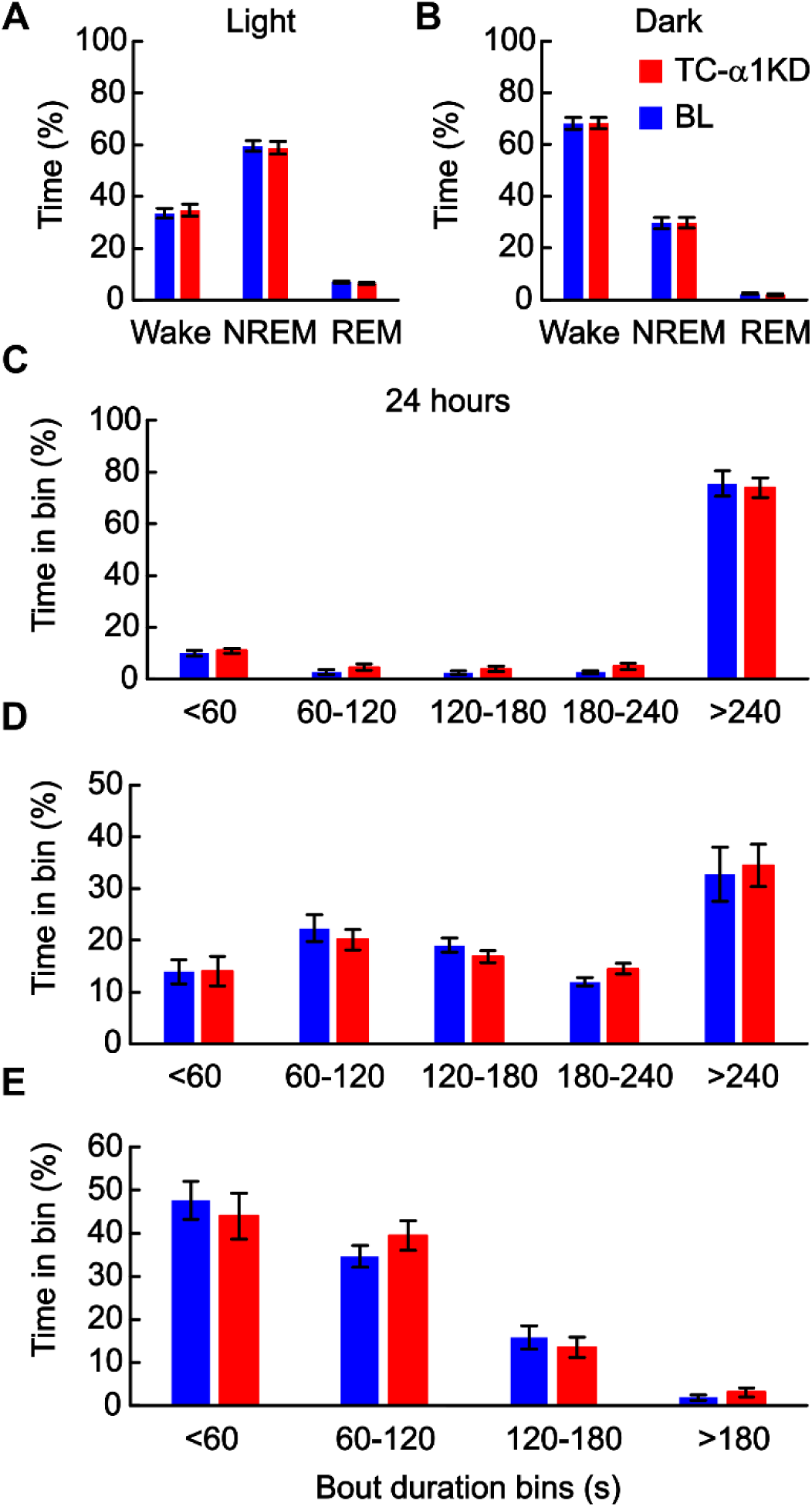
TC-α1KD does not disrupt normal sleep-wake architecture in mice. **A)** Compared to their baseline (BL) records (blue), TC-α1KD (red) mice had no detectable alterations to their proportions of sleep-wake states during the 12h light period. **B)** Compared to their BL records (blue), TC-α1KD (red) mice had no detectable alterations to their proportions of sleep-wake states during the 12h dark period. **C)** Compared to their BL records (blue), TC-α1KD (red) mice had no detectable alterations to their wake bout profiles, as determined by time-weighted bout analysis over 24h. **D)** Compared to their BL records (blue), TC-α1KD (red) mice had no detectable alterations to their NREM sleep bout profiles, as determined by time-weighted bout analysis over 24h. **E)** Compared to their BL records (blue), TC-α1KD (red) mice had no detectable alterations to their REM sleep bout profiles, as determined by time-weighted bout analysis over 24h. N = 14.

**Fig S2.**
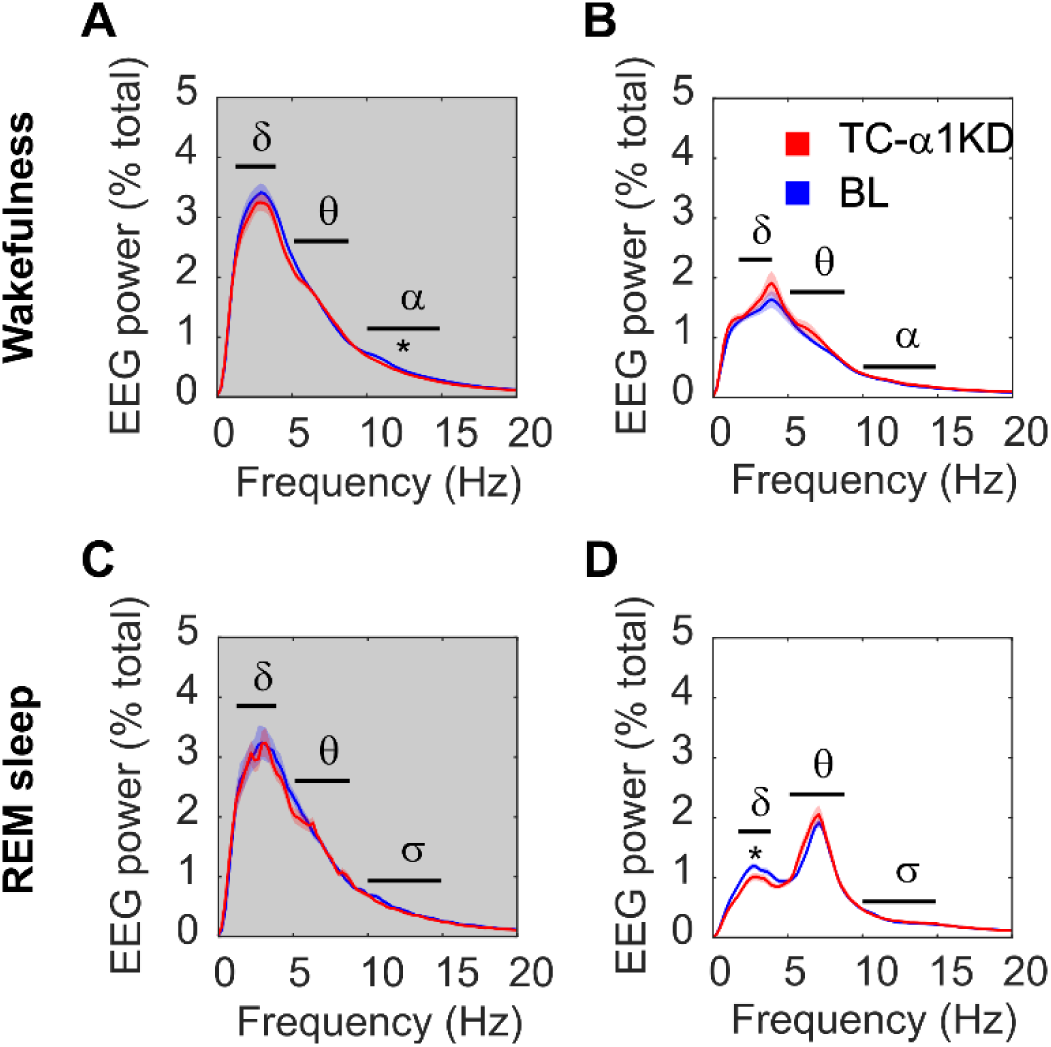
Power spectral density of wakefulness and REM sleep show inconsistent changes to alpha and delta power following TC-α1KD. **A)** Waking delta (δ) and theta (θ) power were not different between baseline (BL) and TC-α1KD records, during the 12h dark period, alpha (α) power was significantly reduced in TC-α1KD mice compared with their BL records. **B)** Waking delta (δ) theta (θ) and alpha (α) power were not different between BL and TC-α1KD records, during 12h the light period. **C)** REM sleep delta (δ) theta (θ) and sigma (σ) power were not different between BL and TC-α1KD records, during 12h the dark period. **D)** Waking delta (δ) power was significantly reduced in TC-α1KD mice compared with their baseline records, theta (θ) and sigma (σ) power were not different between BL and TC-α1KD records, during the 12h light period. Significance was determined using two-tailed paired *t*-tests. Thick colored lines indicate mean; envelopes indicate SEM. Thick black lines indicate areas tested for significance. N = 14.

**Fig S3.**
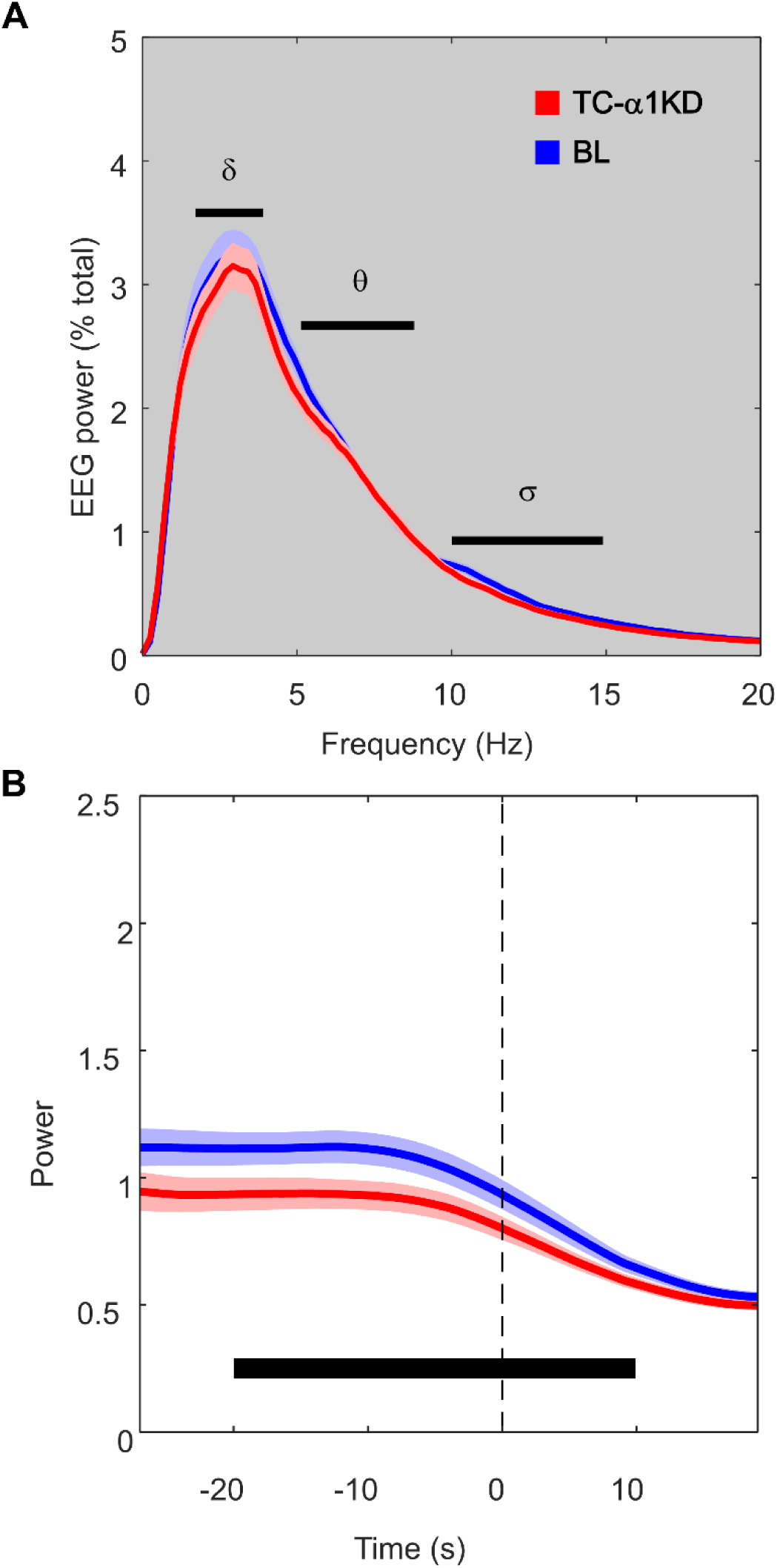
Sigma power is consistently attenuated during NREM following TC-α1KD. **A)** NREM sigma (σ) power was significantly reduced in TC-α1KD mice compared with their baseline (BL) records. Delta (δ) and theta (θ) power were not different between BL and TC-α1KD records, during the 12h dark period, similar to the findings in the light period shown in main figure 1C. **B)** Compared with their BL levels (blue), TC-α1KD mice (red) had reduced sigma (σ; 10 – 15 Hz) power in NREM immediately before a transition to wakefulness [t *(13)* = - 2.9, *p* = 0.012], similar to the effect shown in NREM-REM transitions in the main figure 1F. Significance was tested using a two-tailed paired *t*-test. Thick colored lines indicate mean; envelopes indicate SEM. The thick black bar indicates the area that was tested for significance. N = 14.

**Supp Table 1.** The number of times a thalamic nucleus was identified as transduced by AAVs delivering sgRNAs against its target GABA_A_ receptor subunit. Counts were performed manually by visual comparison with a mouse brain atlas.

| Thalamic Nucleus | TC-α1KD repeated measures group | TC-α1KD fear conditioning group | TC-α3KD fear conditioning group |
| --- | --- | --- | --- |
| Paraventricular | 10 | 6 | 6 |
| Mediodorsal | 11 | 5 | 6 |
| Anteriodorsal | 9 | 3 | 0 |
| Anteromedial | 8 | 0 | 2 |
| Paratenial | 8 | 5 | 6 |
| Central Medial | 6 | 4 | 6 |
| Interanteromedial | 6 | 0 | 3 |
| Reuniens | 4 | 0 | 0 |
| Ventral Reuniens | 4 | 0 | 0 |
| Xiphoid | 3 | 0 | 0 |
| Paraxiphoid | 1 | 0 | 0 |

## Notes

### Summary of Updates

Formatting revisions and edits were made to target specific journals.

