## Supplemental Materials for "Synaptic GABA dysfunction of thalamocortical neurons impairs sleep spindle morphology and fear extinction"

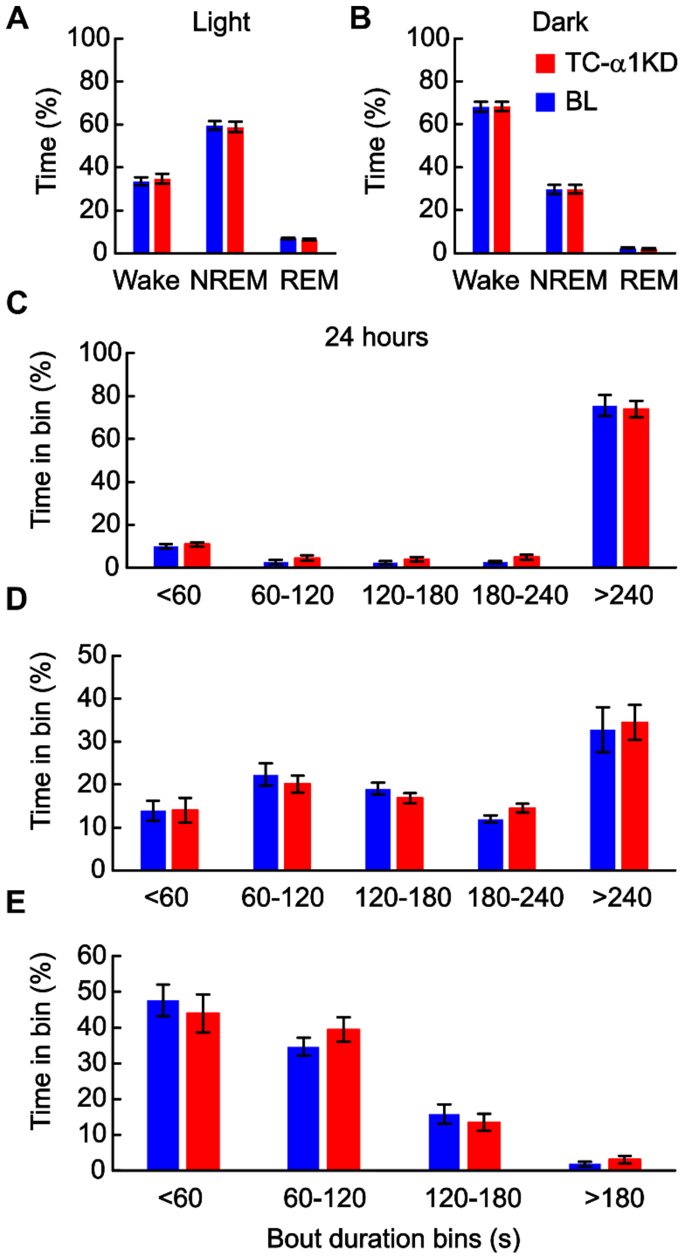


**Fig S1. TC-α1KD does not disrupt normal sleep-wake architecture in mice. A)** Compared to their baseline (BL) records (blue), TC-α1KD (red) mice had no detectable alterations to their proportions of sleep-wake states during the 12h light period. **B)** Compared to their BL records (blue), TC-α1KD (red) mice had no detectable alterations to their proportions of sleep-wake states during the 12h dark period. **C)** Compared to their BL records (blue), TC-α1KD (red) mice had no detectable alterations to their wake bout profiles, as determined by time-weighted bout analysis over 24h. **D)** Compared to their BL records (blue), TC-α1KD (red) mice had no detectable alterations to their NREM sleep bout profiles, as determined by time-weighted bout analysis over 24h. **E)** Compared to their BL records (blue), TC-α1KD (red) mice had no detectable alterations to their REM sleep bout profiles, as determined by time-weighted bout analysis over 24h. N = 14.

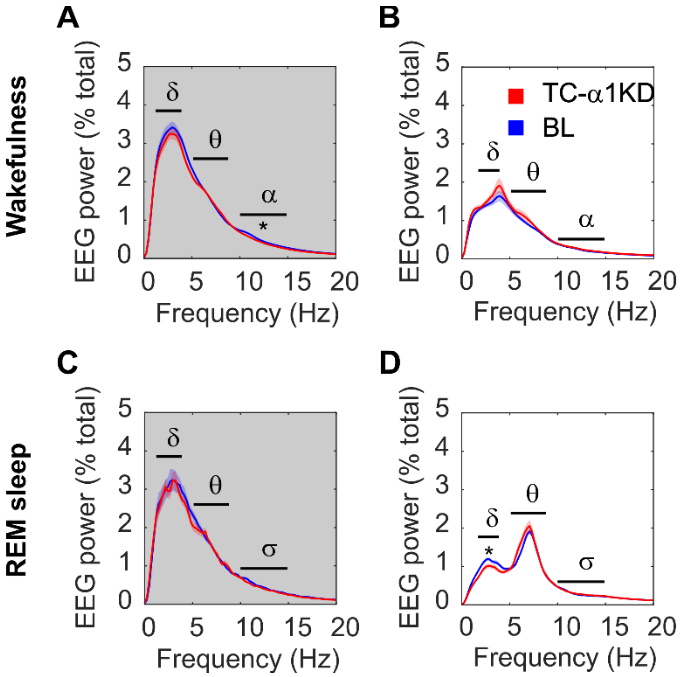


**Fig S2. Power spectral density of wakefulness and REM sleep show inconsistent changes to alpha and delta power following TC-α1KD. A)** Waking delta (δ) and theta (θ) power were not different between baseline (BL) and TC-α1KD records, during the 12h dark period, alpha (α) power was significantly reduced in TC-α1KD mice compared with their BL records. **B)** Waking delta (δ) theta (θ) and alpha (α) power were not different between BL and TC-α1KD records, during 12h the light period. **C)** REM sleep delta (δ) theta (θ) and sigma (σ) power were not different between BL and TC-a1KD records, during 12h the dark period. **D)** Waking delta (δ) power was significantly reduced in TC-α1KD mice compared with their baseline records, theta (θ) and sigma (σ) power were not different between BL and TC-α1KD records, during the 12h light period. Significance was determined using two-tailed paired *t*-tests. Thick colored lines indicate mean; envelopes indicate SEM. Thick black lines indicate areas tested for significance. N = 14.

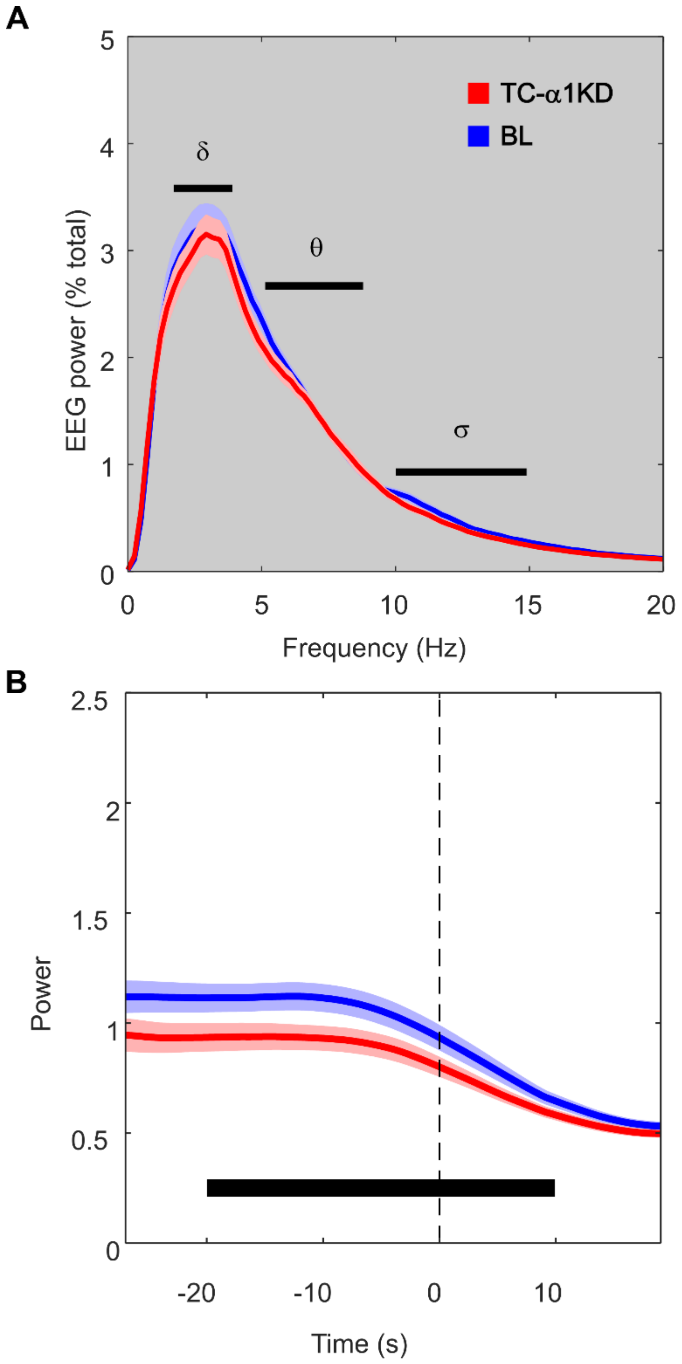


**Fig S3. Sigma power is consistently attenuated during NREM following TC-α1KD.** **A)** NREM sigma (σ) power was significantly reduced in TC-α1KD mice compared with their baseline (BL) records. Delta (δ) and theta (θ) power were not different between BL and TC-α1KD records, during the 12h dark period, similar to the findings in the light period shown in main figure 1C. **B)** Compared with their BL levels (blue), TC-α1KD mice (red) had reduced sigma (σ; 10 – 15 Hz) power in NREM immediately before a transition to wakefulness [t *(13)* = -2.9, *p* = 0.012], similar to the effect shown in NREM-REM transitions in the main figure 1F. Significance was tested using a two-tailed paired *t*-test. Thick colored lines indicate mean; envelopes indicate SEM. The thick black bar indicates the area that was tested for significance. N = 14.

| Thalamic Nucleus | $\alpha$1KD repeated measures group | $\alpha$1KD fear conditioning group | $\alpha$3KD fear conditioning group |
|---|---|---|---|
| Paraventricular | 10 | 6 | 6 |
| Mediodorsal | 11 | 5 | 6 |
| Stria medularis | 11 | 4 | 5 |
| Anteriodorsal | 9 | 3 | 0 |
| Anteriomedial | 8 | 0 | 2 |
| Paratenial | 8 | 5 | 6 |
| Central Medial | 6 | 4 | 6 |
| Interanteromedial | 6 | 0 | 3 |
| Reuniens | 4 | 0 | 0 |
| Ventral Reuniens | 4 | 0 | 0 |
| Xiphoid | 3 | 0 | 0 |
| Paraxiphiod | 1 | 0 | 0 |

Supp Table 1. The number of times a thalamic nucleus was identified as transduced by AAVs delivering sgRNAs against its target $GABA_A$ receptor subunit. Counts were performed manually by visual comparison with a mouse brain atlas.
